# Clustered Cases of Human Adenovirus types 4, 7 and 14 Infections during the 2018 - 2019 Season revealed by US Department of Defense Respiratory Pathogen Surveillance and Whole Genome Sequencing

**DOI:** 10.1101/2022.11.11.516166

**Authors:** Adam R. Pollio, Anthony C. Fries, Yu Yang, Jerry J. Hughes, Christian K. Fung, Matthew A. Conte, Robert A. Kuschner, Natalie D. Collins, Elizabeth A. Macias, Jun Hang

## Abstract

Human adenoviruses (HAdV) are genetically diverse and can infect a number of tissues with severities varied from mild to fatal. HAdV types 3, 4, 7, 11, 14, 21 and 55 were associated with acute respiratory illnesses (ARI) outbreaks in the US and in other countries. The risk of outbreaks can be effectively controlled by HAdV vaccination or mitigated by screening and preventive measures. During the influenza season 2018 – 2019, the DoD Global Respiratory Pathogen Surveillance Program (DoDGRS) received 24,300 specimens. HAdV samples that produced positive cytopathic effects in viral cultivation were subjected to next-generation sequencing for genome sequence assembly, genome typing, whole genome phylogeny, and sequence comparative analyses. Variety of HAdV types were identified in this study, including HAdV types 1 – 7, 14, 55 and 56. HAdV types 4, 7 and 14 were found in clustered cases in Colorado, Florida, New York, and South Carolina. Comparative sequence analyses of these isolates revealed the emergence of novel genetic mutations despite the stability of adenovirus genomes. Genomic surveillance of HAdV related to possible outbreaks, shed light on prevalence, genetic divergence, and viral evolution of human adenoviruses. Continued surveillance will inform risk assessment and countermeasures.

## Introduction

Human adenoviruses (family *Adenoviridae*, genus *Mastadenovirus*) (HAdV) are common viral pathogens that cause multiple respiratory, gastrointestinal and eye infections (1, 2). Symptoms in healthy individuals are often mild and self-limited but can be severe and even fatal in patients with underdeveloped, compromised or suppressed immunity (3). However, diseases associated with HAdVs, in particular, the rate of acute respiratory illnesses (ARI) caused by respiratory HAdVs, are believed to be underreported in the general population. Respiratory HAdV types 3, 4, 7, 11, 14, 21 and the more recent type 55 have been reported causes of ARI outbreaks in the US and in other countries (1, 4–6). HAdV ARI outbreaks in the US were documented in case reports and retrospective and prospective analyses (4, 6–9). Active surveillance on respiratory HAdV infection with detailed characterization is desired to inform mitigation measures to protect high risk populations (10). The US military has successfully used licensed HAdV vaccines to prevent ARI outbreak during recruit basic training, and experienced persistent ARI outbreaks when HAdV vaccines were not used (11, 12). The US Department of Defense Global Emerging Infection Surveillance (DoD GEIS) program actively monitors infectious pathogens including respiratory viruses, not just in military recruits, and includes DoD beneficiaries (13). In this study, we conducted genomic surveillance for HAdVs and report the finding of HAdV types 4, 7 and 14 that caused clustered cases in active duty members and their dependents during the season 2018 – 2019. Whole genome comparative analysis revealed novel genetic mutation, but overall conservation of these adenovirus genomes.

## Materials and methods

The HAdV isolates were obtained from the DoD Global Respiratory Pathogen Surveillance Program (DoDGRPSP) at the US Air Force School of Aerospace Medicine (USAFSAM), Wright-Patterson AFB, Ohio. The case definition, sample collection and the season 2018 – 2019 surveillance were described by GD. Kersellius, et al (14). The respiratory specimens positive for HAdV in molecular assay were subjected to viral culture using cell lines A549, RM2, etc. The supernatant from cultures with positive cytopathic effect (CPE) were extracted to purify HAdV genomic DNA for whole genome sequencing using QIAseq FX DNA Library Kit (QIAGEN, https://www.qiagen.com) and MiSeq next-generation sequencing (NGS) system and Reagent Kit v3 (600-cycle) (Illumina, https://www.illumina.com) at the Walter Reed Army Institute of Research (WRAIR). The bioinformatics tools, NGS_Mapper, Geneious R10 (Biomatters Ltd., https://www.geneious.com), Integrative Genomics Viewer (IGV) (Broad Institute, https://igv.org), were used for whole genome sequence assembly and visualization. Multiple sequence alignments were generated using MUSCLE and phylogenetic trees built using Neighbor-Joining method and Tamura-Nei genetic distance model, with 500 bootstrap replicates, as described previously (5). Representative complete genome sequences for HAdV types 4, 7 and 14 strains from different countries and collection years were retrieved from GenBank and included in whole genome phylogenetic analyses. Sequences containing large segment recombinant, such as HAdV7 strains HAdV 7/Haiti-0707/2014 (MN531562) and HAdV-B/USA/8010/1998 (MH910665), were excluded.

## Results

DoDGRPSP detected 320 HAdVs (1.3% of all cases, or 2.8% of all cases with one or multiple pathogens detected) in the season 2018 – 2019 (14). Among the total of 161 HAdV isolates available for whole genome sequencing, a variety of types were identified in the US and other countries, including HAdV-1 (N = 15), HAdV-2 (N = 26), HAdV-3 (N = 60), HAdV-4 (N = 13), HAdV-5 (N = 6), HAdV-6 (N = 3), HAdV-7 (N = 12), HAdV-14 (N = 10), HAdV-55 (N = 1), and HAdV-56 (N = 1) (**Supplemental Figure 1**). The HAdV type distribution is consistent with the 2003-2016 and 2017 HAdV surveillance data from CDC’s National Adenovirus Type Reporting System (NATRS) (https://www.cdc.gov/adenovirus/reporting-surveillance/natrs/surveillance-data.html), with Pearson’s correlation p-value of 7.27E-04 and 1.36E-03, respectively. HAdV types 1-3 were seen in multiple states and different times (**Supplemental Figure S1**). In contrast, the HAdV-4, HAdV-7 and HAdV14 were identified as geographically clustered cases (**Table 1, Figure 1**, **2** and **S1**), suggesting possible outbreaks. The 49 cases consist of 39 (79.6%) males and 10 (20.4%) females. All patients were young, with age group (number of cases) as follows, 0-5 (5), 6-10 (3), 11-15 (2), 16-20 (23), 21-25 (13), 26-30 (2) and 31-35 (1). None of these cases were reported having pneumonia, one individual was hospitalized. The assembled complete genome sequences for HAdV-4, 7, and 14 were subjected to further manual curation, full annotations, GenBank submission (accession numbers shown in **Table 1**) and phylogenetic and comparative sequence analyses.

**Table 1.** Human adenovirus types 4, 7 and 14 isolates from this study.

| **HAdV Strain** | **Type** | **Collection Time** | **Collection Site** | **Age Bin** | **Gender** | **GenBank Accession** |
| --- | --- | --- | --- | --- | --- | --- |
| 21660 S4TF1 | 4 | 7-Dec-18 | New York | 16-20 | Male | OM649219 |
| 21662 S4TF3 | 4 | 25-Feb-19 | New York | 16-20 | Female | OM649220 |
| 21665 S4TF6 | 4 | 11-Feb-19 | New York | 16-20 | Male | OM649224 |
| 21666 S4TF7 | 4 | 7-Mar-19 | New York | 16-20 | Male | OM649227 |
| 21667 S4TF8 | 4 | 18-Mar-19 | New York | 16-20 | Male | OM649229 |
| 21668 S4TF9 | 4 | 18-Mar-19 | New York | 16-20 | Male | OM649230 |
| 21669 S4TF10 | 4 | 6-Mar-19 | New York | 16-20 | Male | OM649228 |
| 21671 S4TF12 | 4 | 5-Feb-19 | New York | 21-25 | Male | OM649221 |
| 21672 S4TF13 | 4 | 18-Mar-19 | New York | 21-25 | Male | OM649223 |
| 21673 S4TF14 | 4 | 20-Mar-19 | New York | 21-25 | Male | OM649226 |
| 21677 S4TF18 | 4 | 6-Apr-19 | New York | 21-25 | Male | OM649222 |
| 21678 S4TF19 | 4 | 12-Feb-19 | New York | 21-25 | Male | OM649231 |
| 21679 S4TF20 | 4 | 14-Feb-19 | New York | 21-25 | Male | OM649225 |
| 21701 S4TF42 | 4 | 27-Nov-18 | Virginia | 0-5 | Male | OM649218 |
| 21966 S4TF57 | 4 | 18-Mar-19 | Colorado | 16-20 | Male | OM649216 |
| 21967 S4TF58 | 4 | 5-Mar-19 | Colorado | 21-25 | Male | OM649214 |
| 21968 S4TF59 | 4 | 26-Sep-18 | Colorado | 16-20 | Female | OM649209 |
| 21969 S4TF60 | 4 | 28-Sep-18 | Colorado | 16-20 | Male | OM649205 |
| 21970 S4TF61 | 4 | 24-Oct-18 | Colorado | 16-20 | Male | OM649207 |
| 21972 S4TF63 | 4 | 4-Mar-19 | Colorado | 16-20 | Male | OM649212 |
| 21973 S4TF64 | 4 | 20-Mar-19 | Colorado | 16-20 | Male | OM649213 |
| 21974 S4TF65 | 4 | 10-Oct-18 | Colorado | 16-20 | Male | OM649208 |
| 21975 S4TF66 | 4 | 28-Sep-18 | Colorado | 21-25 | Male | OM649210 |
| 21976 S4TF67 | 4 | 15-Mar-19 | Colorado | 21-25 | Male | OM649215 |
| 21984 S4TF75 | 4 | 30-Nov-18 | Florida | 6-10 | Female | OM649206 |
| 21947 S4TF137 | 4 | 30-Jan-19 | Ohio | 16-20 | Female | OM649211 |
| 22128 S4TF161 | 4 | 13-Nov-18 | Texas | 11-15 | Female | OM649217 |
| 21674 S4TF15 | 7 | 14-Dec-18 | New York | 21-25 | Male | OM453826 |
| 21675 S4TF16 | 7 | 30-Mar-19 | New York | 21-25 | Male | OM453822 |
| 21676 S4TF17 | 7 | 2-Apr-19 | New York | 21-25 | Male | OM453825 |
| 21680 S4TF21 | 7 | 27-Sep-18 | Florida | 0-5 | Female | OM453830 |
| 21682 S4TF23 | 7 | 3-Nov-18 | Florida | 0-5 | Male | OM453832 |
| 21703 S4TF44 | 7 | 14-Nov-18 | Florida | 31-35 | Male | OM453831 |
| 21971 S4TF62 | 7 | 25-Jan-19 | Colorado | 16-20 | Male | OM453823 |
| 21987 S4TF78 | 7 | 15-Oct-18 | Florida | 0-5 | Male | OM453829 |
| 21993 S4TF84 | 7 | 30-Oct-18 | South Dakota | 0-5 | Female | OM453824 |
| 21939 S4TF129 | 7 | 10-Jan-19 | North Carolina | 6-10 | Male | OM453827 |
| 22124 S4TF157 | 7 | 19-Nov-18 | New Jersey | 11-15 | Male | OM453828 |
| 22127 S4TF160 | 7 | 29-Nov-18 | Florida | 6-10 | Female | OM453833 |
| 22113 S4TF146 | 14 | 21-Nov-18 | South Carolina | 16-20 | Female | OM514959 |
| 22114 S4TF147 | 14 | 2-Jan-19 | South Carolina | 21-25 | Male | OM514964 |
| 22115 S4TF148 | 14 | 6-Nov-18 | South Carolina | 26-30 | Male | OM514960 |
| 22117 S4TF150 | 14 | 16-Jan-19 | South Carolina | 16-20 | Male | OM514965 |
| 22118 S4TF151 | 14 | 1-Apr-19 | South Carolina | 16-20 | Male | OM514962 |
| 22119 S4TF152 | 14 | 31-Mar-19 | South Carolina | 16-20 | Male | OM514963 |
| 22120 S4TF153 | 14 | 9-Jan-19 | South Carolina | 16-20 | Male | OM514966 |
| 22121 S4TF154 | 14 | 27-Mar-18 | South Carolina | 16-20 | Male | OM514961 |
| 22122 S4TF155 | 14 | 28-Mar-19 | South Carolina | 16-20 | Male | OM514958 |
| 21125 S4TF158 | 14 | 6-Nov-18 | South Carolina | 26-30 | Female | OM514957 |

**Figure 1.**
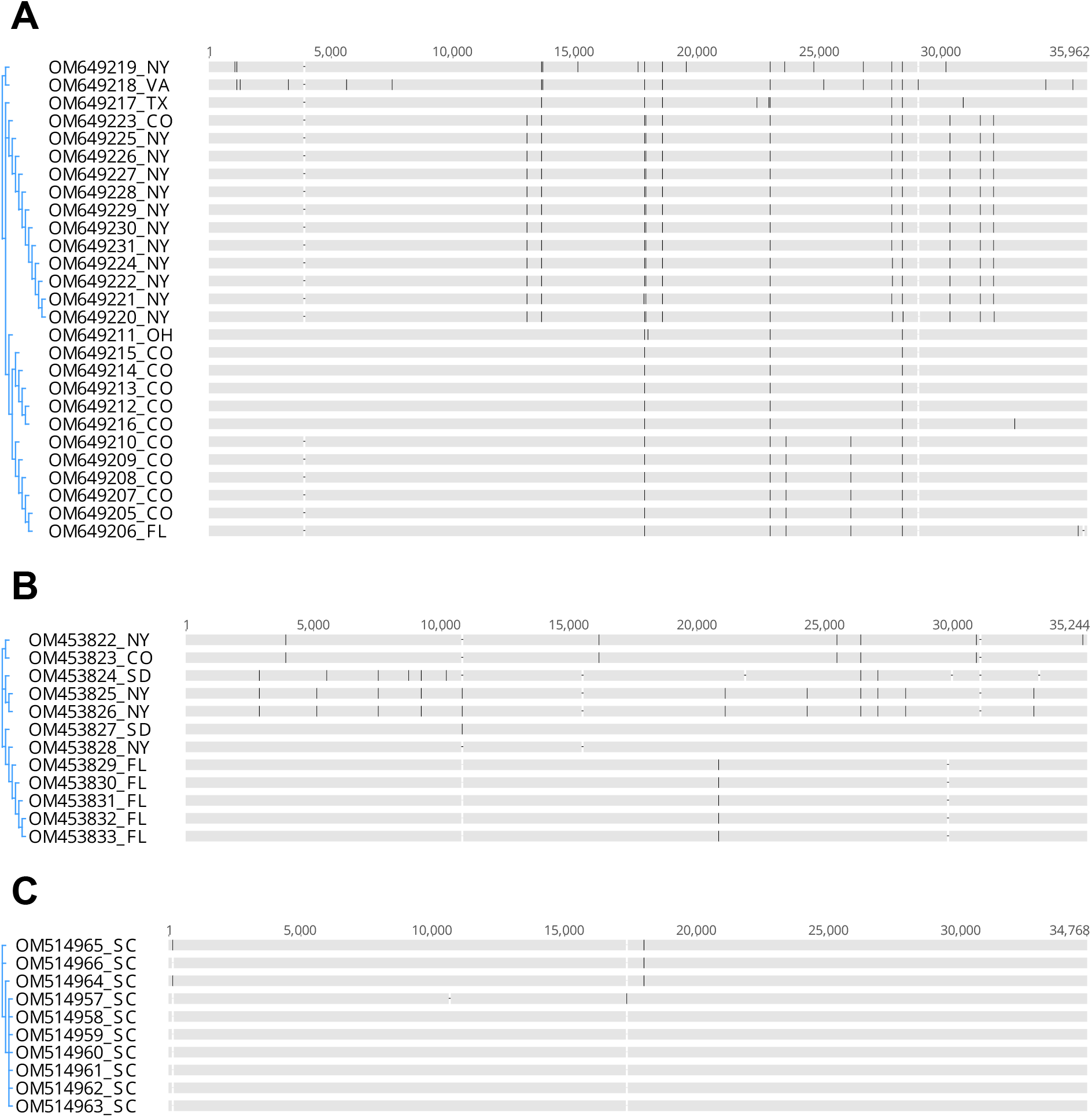
Multiple sequence alignment of complete genome sequences of human adenoviruses (HAdV) of season 2018 – 2019 in this study. **(A)** HAdV 4; **(B)** HAdV 7; **(C)** HAdV 14. Genomes are arranged based on phylogenetic relatedness.

**Figure 2.**
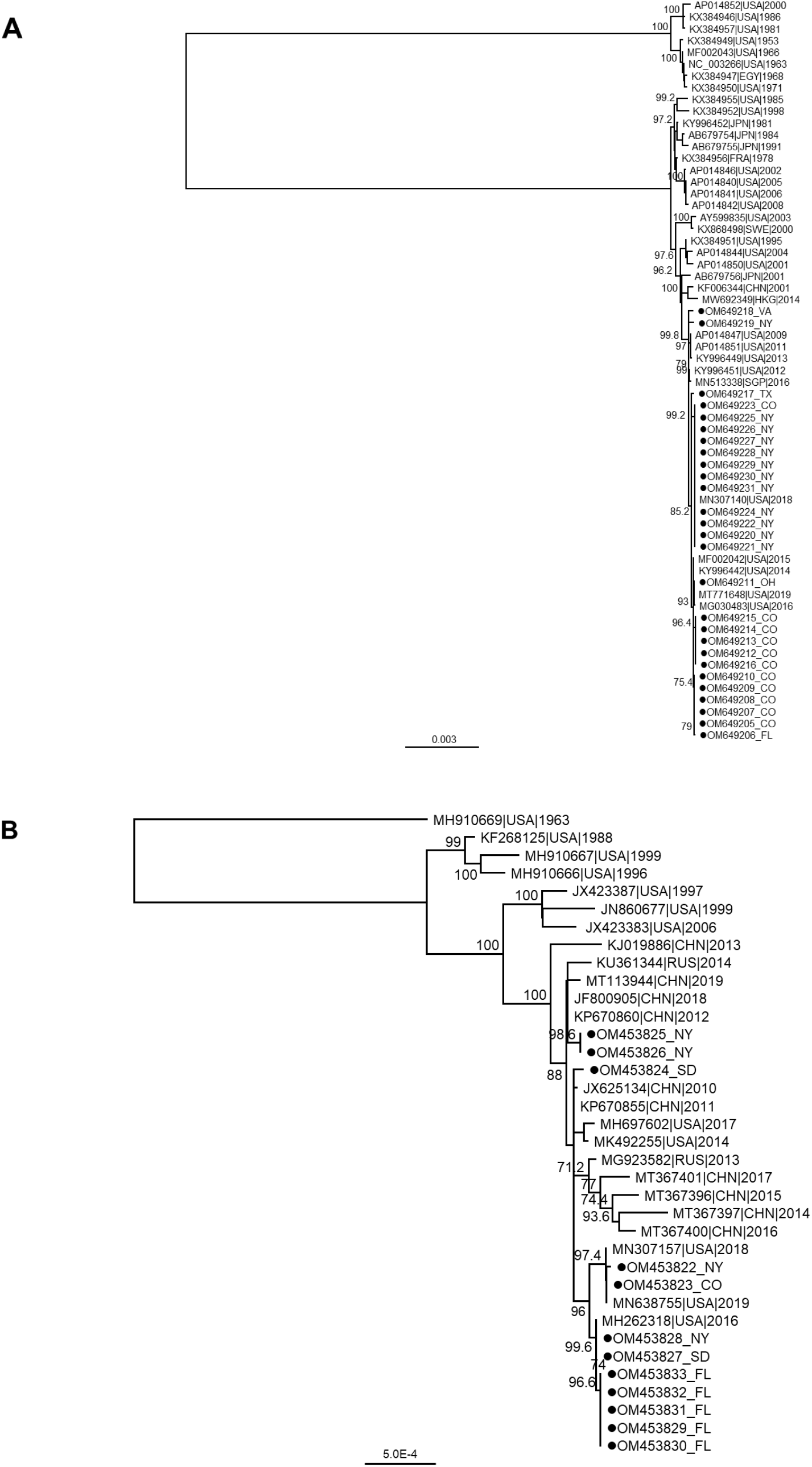
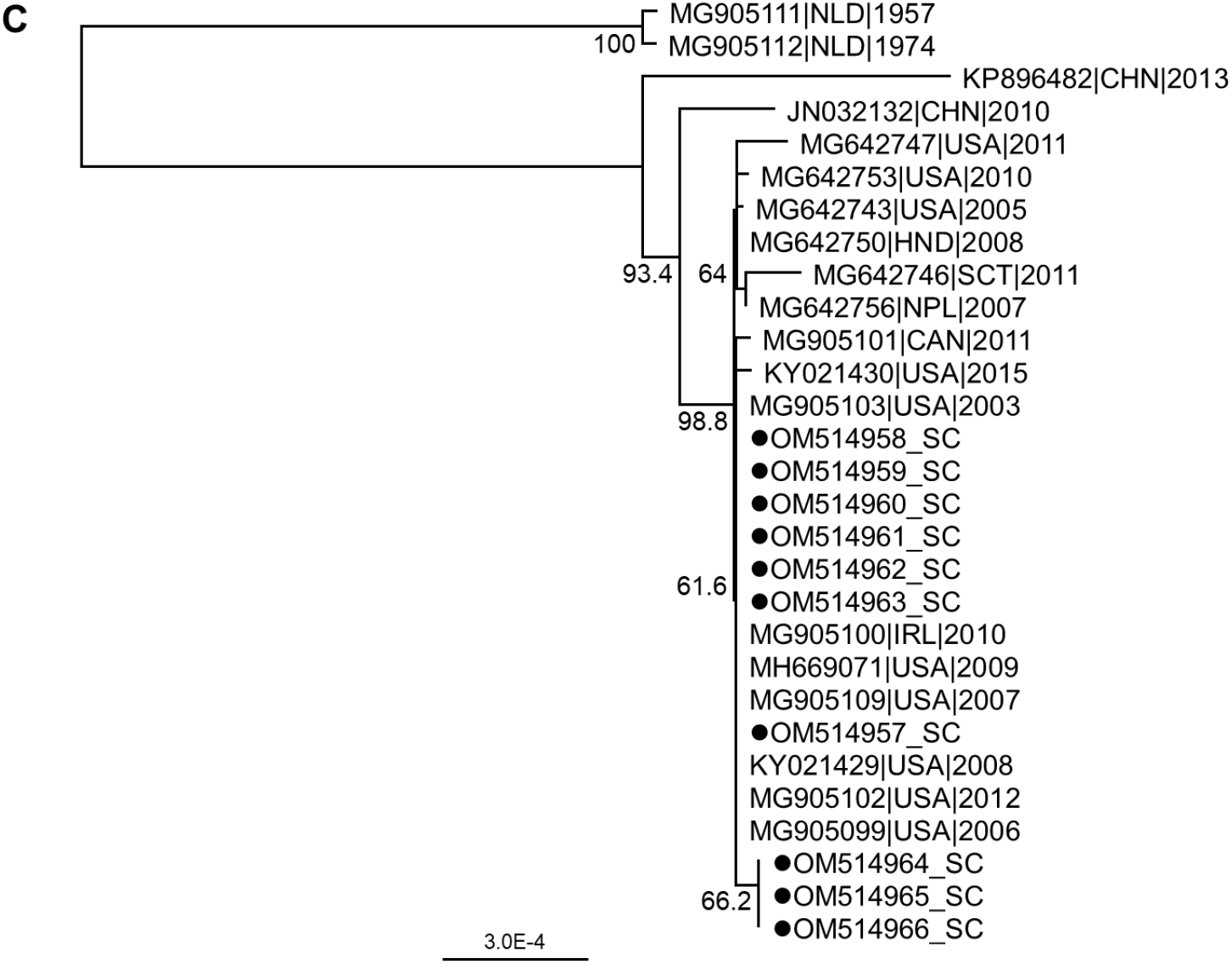
Whole genome phylogeny of human adenoviruses (HAdV) in this study and representative HAdV sequences in GenBank. **(A)** HAdV 4; **(B)** HAdV 7; **(C)** HAdV 14. Neighbor-Joining phylogenetic tree with bootstrap support values from 500 replicates shown at the branches. The scale bar represents estimated nucleotide substitutions per site. The dots indicate the HAdVs from season 2018-2019 in this study.

The HAdV-14 isolates were all from cases in South Carolina. The HAdV-14 genome sequences are nearly identical with each other, with few nucleotide differences in untranslated regions and one synonymous substitution C18006T corresponding to amino acid residue Valine 202 in protein VI. The three genomes carrying the substitution C18006T are from the cases with collection dates separated by about a week, suggesting a potential transmission of HAdV-14 and cluster of cases. Overall, we have seen high sequence similarity shared among HAdV-14 sequences from this study and known sequences of other recent strains in US and worldwide, with less than 0.4% of nucleotide differences from the prototype strain Netherlands/DeWit/1957 (MG905111).

The sequence alignment and phylogenetic analysis for the 12 HAdV-7 isolates demonstrate the circulation of multiple strains in the US. The five cases in Florida during a period of two months, including one male adult and four young children of different ages, shared identical genome sequences and suggested clustered cases (**Table 1, Figure S1**). Nucleotide differences are present in untranslated regions in between protein coding sequences (CDS), synonymous and nonsynonymous mutations in multiple proteins, including DNA polymerase, terminal protein precursor, hexon, etc. (**Table 2**). The Florida HAdV-7 isolate has a large deletion of 30 nucleotides, corresponding to a 10 amino acid deletion, which is not present in other strains from this study or HAdV-7 sequences in GenBank. This deletion is located in the middle of the 41 amino acid membrane glycoprotein E3 CR 1-delta, which is one of the CR1 proteins that play a complex role in modulating host immune response to viral infection.

**Table 2.** Sequence variations among human adenovirus types 4, 7 and 14 isolates from this study. GenBank sequences KY996443 and KP670860 were used as references to determine position of the nucleotide mutations for human adenovirus types 4 and 7 respectively. There is no amino acid sequence difference among type 14 isolates from this study.

| Gene/Protein | Type | Strains | Amino Acid change | Nucleotide mutation position |
| --- | --- | --- | --- | --- |
| E1B large T antigen | 4 | 21701 | D468N | 3271 |
| DNA polymerase | 4 | 21701 | A371T, R992H | 7506, 5642 |
| Capsid protein precursor pVI | 4 | 21966, 21967, 21972, 21973, 21976 | T175M | 17833 |
|  |  | 21662, 21665, 21666, 21667, 21668, 21669, 21672, 21673, 21677, 21678, 21679 | P199S | 17904 |
|  |  | 21947 | P223S | 17976 |
| Hexon protein | 4 | 21660 | V461I | 19555 |
| DNA-binding protein | 4 | 22128 | E104G | 22454 |
| E3 19 kDa protein | 4 | 21947, 21966, 21967, 21968, 21969, 21970, 21972, 21973, 21974, 21975, 21976, 21984 | N63D | 27978 |
| E3 24.8 kDa protein | 4 | 21966, 21967, 21972, 21973, 21976 | D21E | 28408 |
| Fiber protein | 4 | 21662, 21665, 21666, 21667, 21668, 21669, 21671, 21672, 21673, 21677, 21678, 21679 | E41Q | 31579 |
| 3' inverted terminal repeat (ITR) | 4 | 21984 | 18 bp deletion | 35828 - 35845 |
| Maturation protein IVa2 | 7 | 21674, 21676 | G55R | 5133 |
| DNA polymerase | 7 | 21993 | D1039E | 5517 |
| E2B precursor terminal protein | 7 | 21674, 21676 | K398E | 9208 |
|  |  | 21993 | P68L, K398E | 10197, 9208 |
| Hexon protein | 7 | 21680, 21682, 21703, 21987, 22127 | M818I | 20848 |
| Capsid protein precursor pVIII | 7 | 21674, 21676, 21993 | Q131R | 27063 |
| Membrane glycoprotein E3 CR1-delta | 7 | 21680, 21682, 21703, 21987, 22127 | Δ18-27 | 29787 - 29816 |
| E4 34 kDa protein | 7 | 21674, 21676 | Q87P | 33156 |
| | | 21993 | Δ16-19 | 33361 - 33372 |

Among the 27 HAdV-4 isolates, several strains or variants were identified. These strains are closely related to US strains of recent years, e.g. HAdVE/USA_New York/1418/2015/P4H4F4 (MF002042). The isolate 21660_S4TF1 (OM649219), which is relatively divergent from other isolates, is nearly identical to HAdV-E/USA_WI/5556/2018/P4H4F4 (MN307142) which was reported to be associated with ARI outbreaks at five US colleges (7), also occurring in the season 2018 – 2019, with nucleotide identities of 35959/35960. The only difference is position 19555 A/G, corresponding to hexon I461V. The subclade of isolate 21947 and five other isolates shares almost identical sequence with viruses that caused outbreaks in US Naval Academy in 2016 (MG030483) and US Coast Guard in 2019 (MT771648) (4, 8). One unique finding in this study is a novel 18 nucleotides long deletion presents in the 3’-terminal inverted terminal repeat (ITR) at position 35927 of HAdV4 isolate 21984_S4TF75 (OM649206), while the 5’-terminal ITR remains unchanged.

Nucleotide divergences and amino acid differences among the HAdV-4 and 7 strains from this study were found in a variety of proteins, including DNA polymerase, major and minor capsid proteins hexon, fiber, pVI and pVIII, core proteins DNA terminal protein and IVa, and proteins involved in modulation of host immune responses (**Table 2**). Most of the recently designated HAdV types (e.g. HAdV-55) are genetic recombinants of penton base, hexon, and fiber protein genes. The recombination can lead to emerging infection outbreaks, as shown for HAdV-55 outbreaks in Asia and suggests that this type needs to be monitored (15, 16). The multiple sequence alignments and phylogenetic analyses showed that the HAdV-4, 7 and 14 isolates in this study did not have significant recombination (**Figure 2**).

## Discussion

A variety of HAdV types were identified in this study, suggesting the concurrent circulation of HAdV types and genetic variants in the US. The DoDGRPSP and whole genome sequence analysis in this study enable comprehensive surveillance of respiratory HAdV infections. DoDGRPSP, which mainly covers the US active duty members and their dependents, revealed similar respiratory HAdV prevalence as shown in CDC NATRS system (2). Both systems are passive surveillance based on sample or data voluntarily contributed by participating laboratories. It is important to continue these efforts to accumulate HAdV type data to inform the clinical and public health for timely prevention and control of HAdV-associated ARI outbreak. The incidents, in which HAdV outbreaks led to death of immunocompromised individuals, highlighted the need for investigation on HAdV transmission risk to people with underlining conditions (3, 12). Moreover, find effective countermeasures to protect those who are at higher risk for infectious diseases and avoid the mortality.

HAdV types 4, 7 and 14 were reported causing ARI in the military (4, 8, 17). This study identified these three adenoviruses in one season, 2018 - 2019. The cases were concentrated in a few US states, suggesting possible clusters of transmissions. The detailed sample information available for further analysis are restricted for human subject privacy protection, therefore, insufficient for resolving the course of infections during the season. Previous studies from others and us showed that HAdV transmissions could last for several months with hundreds of cases (9, 18, 19). In this study, we saw HAdV-4, 7 and 14 samples appeared in several months. It is likely that the number of cases was heavily underestimated due to the mild symptoms. The true prevalence and the scale of possible epidemics in these states would need surveillance that is more intensive.

The live HAdV-4 and 7 vaccine is routinely implemented in the US military basic training to prevent HAdV-caused ARI outbreak (4, 12, 20). The studies evoked the need for expanded use of the vaccine to populations other than the recruits (20, 21). The effective use of the vaccine may not only benefit the military during other congregate events, but also mitigate spread to civilians associated with military personnel. Giving the vaccine to civilian will require additional research for FDA approval. Those immunocompromised individuals, who died to HAdV-7 infection during outbreaks (3, 7), might have survived if the vaccine was available to them. Nevertheless, vaccine efficacy for immunocompromised individuals and the effectiveness of protection from vaccination during outbreak are warranted for investigation.

The whole genome sequencing allowed delineation of genetic diversities, including multiple amino acid mutations and deletions in inverted terminal repeat (ITR) and protein-coding regions (**Table 2**). There are over 110 HAdV types designated by HAdV Working Group (http://hadvwg.gmu.edu). The genetic recombination and mutation can lead to emerging and re-emerging of clinically significant HAdV types, such as HAdV-14, 55 and 7d (3, 6, 22). Considering the broad and persistent circulation of numerous HAdVs, consistent efforts on genomic surveillance and deep analyses are essential for enhancement of HAdV surveillance and informed outbreak management.

## Data availability

Complete genome sequences for HAdV types 4, 7 and 14 from this study were deposited in GenBank with accession numbers listed in **Table 1**.

## Acknowledgments

We thank Irina Maljkovic Berry, Nicos Karasavvas, Richard G. Jarman, Grace M. Lidl, and Leonard N. Binn for their leadership and technical support of the project. We thank James S. Hilaire, Nicole R. Nicholas, Tuan K. Nguyen, and April N. Griggs for their assistance in project management and sample tracking, storage, and retrieval. We thank the Defense Health Agency Immunization Healthcare Division (DHA-IHD), Global Emerging Infections Surveillance and Response System (GEIS), Division of the Armed Forces Health Surveillance Branch.

## Disclaimers

Material has been reviewed by the authors’ respective institutions. There is no objection to its presentation and/or publication. The views expressed here are those of the authors and do not reflect the official policy of the Department of the Army, Department of the Navy, Department of Defense or US Government. This is the work of US government employees and may not be copyrighted (17 USC 105).

---

**Figure S1.**
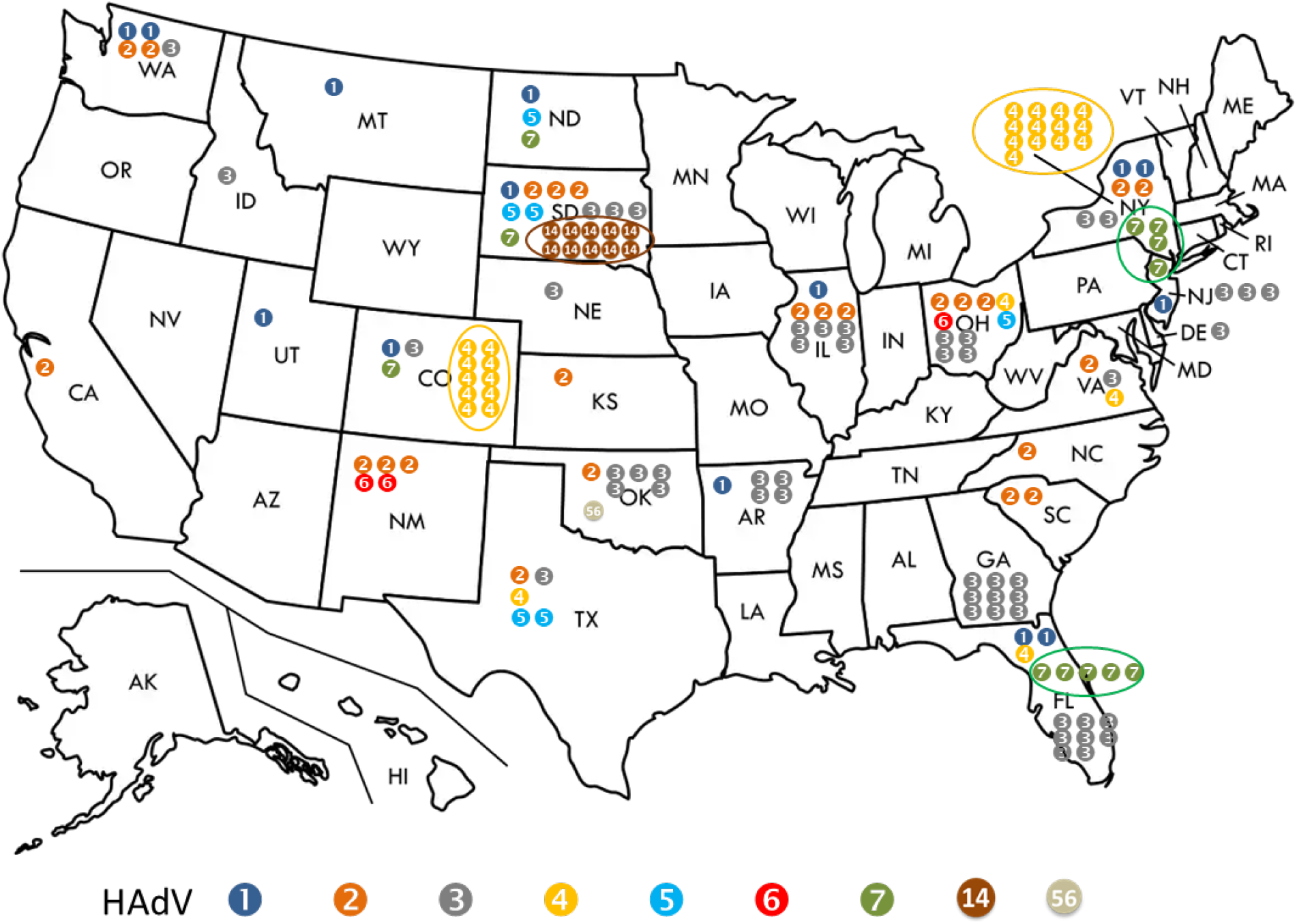
Number and US location of human adenoviruses (HAdV) in this study from season 2018-2019. The dots indicate the HAdV types. The clusters of HAdV-4, 7 and 14 cases are showed in circles. The few HAdVs identified in samples from other countries are not shown, including HAdV-1 from Japan, HAdV-2 from South Korea, HAdV-3 from England (n=7), Italy and Japan, and HAdV-55 from South Korea.

## Notes

### Competing Interest Statement

The authors have declared no competing interest.

