## Supplementary material for "Clustered Cases of Human Adenovirus types 4, 7 and 14 Infections during the 2018 - 2019 Season revealed by US Department of Defense Respiratory Pathogen Surveillance and Whole Genome Sequencing": Figure S1

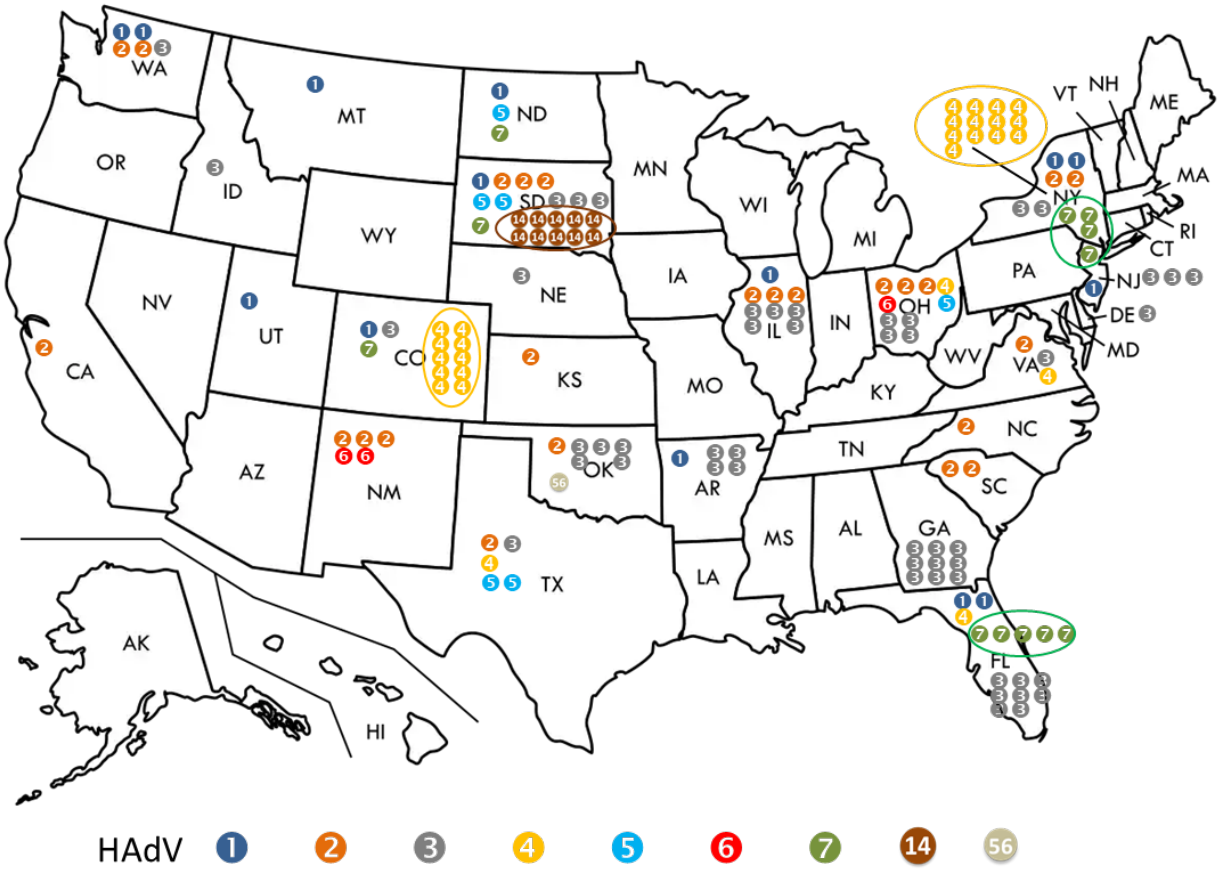


**Figure S1.** Number and US location of human adenoviruses (HAdV) in this study from season 2018-2019. The dots indicate the HAdV types. The clusters of HAdV-4, 7 and 14 cases are showed in circles. The few HAdVs identified in samples from other countries are not shown, including HAdV-1 from Japan, HAdV-2 from South Korea, HAdV-3 from England (n=7), Italy and Japan, and HAdV-55 from South Korea.
